# Heterogeneous dual-metal control of *Salmonella* infection

**DOI:** 10.1101/2023.10.23.562652

**Authors:** Béatrice Roche, Olivier Cunrath, Christopher Bleck, Beatrice Claudi, Minia Antelo Varela, Jiagui Li, Dirk Bumann

## Abstract

Iron controls bacterial infections through diverse pathogen and host mechanisms that remain challenging to disentangle. Here, we determined how individual *Salmonella* cells access iron in infected mice. Our results showed that the iron transporter SLC11A1 restricted iron availability. However, many *Salmonella* bypassed this restriction by targeting macrophage endosomes that contained remnants of iron-rich red blood cells. These iron-replete bacteria dominated overall *Salmonella* growth and masked the relieve of iron-starved bacteria under iron overload. These data, combined with our previous discovery of magnesium deprivation as a primary mechanism for controlling *Salmonella*, reveal a heterogeneous dual-metal mechanism of nutritional immunity, and highlight the power of single-cell analyses under physiological in-vivo conditions to unravel complex anti-bacterial host mechanisms.

**One sentence summary:** Iron and magnesium limitations control distinct *Salmonella* subsets during infection.

## Main text

Iron plays critical roles in bacterial infections as (i) an essential trace nutrient that is restricted by the host (“nutritional immunity”) (*1*); (ii) a cofactor of host antimicrobial enzymes such as NADPH oxidase; (iii) an enhancer of oxidative stress through the Fenton reaction (*2, 3*); and (iv) a regulator of key transcription factors including (NF)-κB, HIF-1α, etc., and IFNγ- and TNFα-signaling (*4-9*). Disentangling the diverse roles of iron remains challenging. For example, in mice infected with invasive *Salmonella*, dietary iron overload exacerbates the infections (*10*), but the interpretation of this observation is complicated. *Salmonella* iron-uptake mutants and reporter strains seem support the idea that *Salmonella’s* replication is iron limitated *(3, 5-7, 9, 11-17)*. However, data on *Salmonella* mutants requiring micromolar iron concentrations might be irrelevant for wild-type *Salmonella* thriving on nanomolar iron (*18*). Iron reporters might yield ambiguous signals because of responses to other stimuli (*19*). Additionally, common bulk-average readouts such as colony-forming units may be confounded by heterogeneous iron levels across tissues (*20*). Lowering iron levels in the host actually increases *Salmonella* growth (*21-24*), contradicting the idea of iron limitation. Therefore, novel approaches are needed to unravel the complex interplay between bacterial and host factors in *Salmonella* infections.

### Divergent *Salmonella* access to iron in mouse spleen

To determine iron access, we developed an iron-responsive *Salmonella* reporter. The *P*_ryhB-2_ *(P*_isrE_) promoter was induced ∼100-fold by iron starvation but did not respond to zinc, manganese, or magnesium starvation (sfig. 1a,b). We fused *P*_ryhB-2_ to genes encoding destabilized variants (*25*) of fluorescent proteins GFP or BFP to monitor current promoter activities. *Salmonella* had low *P*_ryhB-2_ activity in the spleen of commonly used mouse strains (BALB/c, C57BL/6) (Fig. 1C; fig. S2), indicating non-limiting iron access (*18, 26*). These mice are hypersusceptible to *Salmonella* infection (*27*) and have distorted iron homeostasis because they are defective for the metal transporter SLC11A1 (*28-31*).

**Fig. 1.**
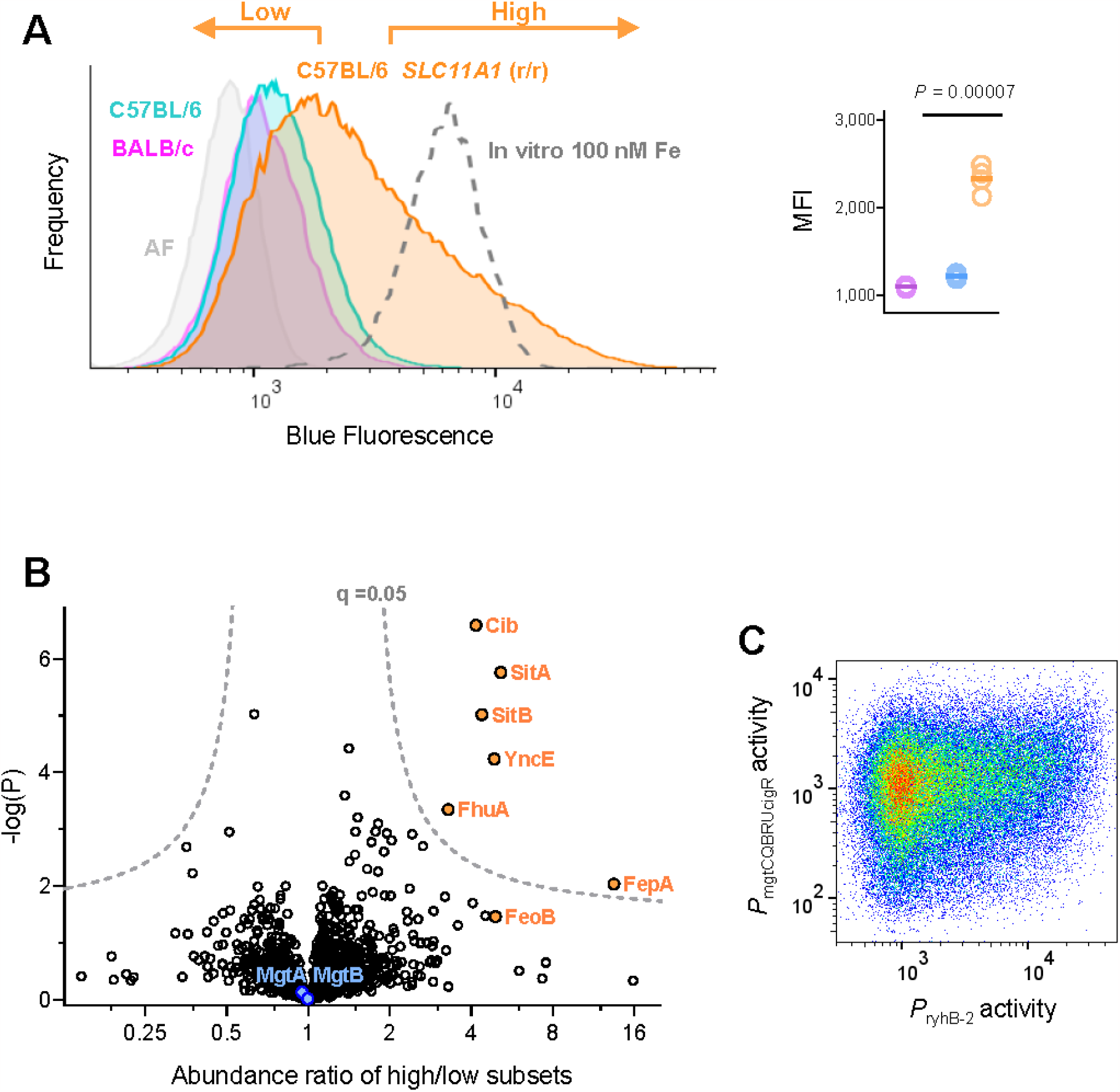
Heterogeneous iron access of *Salmonella* in mouse spleen. **(A)** Blue fluorescence of *Salmonella* carrying *P*_ryhB-2_-*bfp-ova* in spleen of BALB/c, C57Bl/6, or C57Bl/6 mice with restored functional SLC11A1 (C57Bl/6 *SLC11A1*^r/r^) (pooled data for three mice per group; for gating strategy to identify *Salmonella* cells see fig. S1). For comparison, *Salmonella* in-vitro cultures with low iron levels (dashed line) and Salmonella without *bfp* in spleen of C57Bl/6 *SLC11A1*^r/r^ mice (AF, autofluorescence) are also shown. The arrows indicate sorting gates for *Salmonella* subsets. The right graph shows median blue fluorescence intensities (MFI) for individual mice (unpaired t-test). **(B)** Proteome comparison of *Salmonella* with low or high *P*_ryhB-2_-*bfp-ova* activity in C57Bl/6 *SLC11A1*^r/r^ mice (using sorting gates shown in C). Proteins involved in iron uptake are labeled in orange, proteins involved in magnesium uptake are shown in blue. Data for six mice are shown (q, false-discovery rate). **(C)** *P*_ryhB-2_ and *P*_mgtCQBRUcigR_ activities in single *Salmonella* cells in C57Bl/6 *SLC11A1*^r/r^ mice. Pooled data for three independently infected mice are shown. The Spearman rank correlation coefficients for the individual mice were 0.19, 0.17, and 0.19.

In mice with restored functional *SLC11A1* (C57BL/6 *SLC11A1*^r/r^) (*18*), *Salmonella* had higher and more heterogeneous *P*_ryhB-2_ activities (Fig. 1C). We sorted *Salmonella* subsets with low or high *P*_ryhB-2_ activity from spleen using flow cytometry. Mass spectrometry revealed almost identical protein profiles but six out of 1,372 quantified proteins showed significantly higher levels in the *P*_ryhB-2_ high subset (Fig. 1D). These six proteins were involved in iron uptake, consistent with iron starvation of this subset. *P*_ryhB-2_ can respond to other stimuli including oxidative/nitrosative stresses and carbon starvation (*32, 33*). Signature proteins for these stimuli did not differ between *Salmonella* with low or high *P*_ryhB-2_ activity (fig. S3a), suggesting iron starvation as the only relevant cause of heterogeneity. This indicates that some *Salmonella* cells face SLC11A1-mediated iron starvation while other Salmonella cells in the same tissue are iron-replete.

In addition to iron limitation, SLC11A1 also restricts *Salmonella’s* access to magnesium (*18*), suggesting a potential correlation between the availability of these two metals. However, the abundance of the Mg^2+^ transporters MgtA and MgtB was similar in the iron-starved and iron-replete subsets (Fig. 1D). Moreover, there was no correlation between *P*_mgtCQBRUcigR_ (which drives *mgtB* expression and is induced by magnesium starvation) and *P*_ryhB-2_ at the single-cell level (Fig. 1E; fig. S4a,b). Thus, SLC11A1 diminishes overall access of *Salmonella* to both magnesium (29) and iron, but iron starvation in a subset of *Salmonella* is relieved by independent processes.

### Erythrophagocytosis supplies iron to intracellular *Salmonella*

To identify these processes, we determined microenvironments of iron-replete and iron-starved *Salmonella. Salmonella* infected mostly F4/80-positive red pulp macrophages (*34*). Iron-replete *Salmonella* (low *P*_ryhB-2_ activity) resided almost exclusively in macrophages that also contained large (Ø > 2.5 μm) LAMP1/2-positive endosomes, while iron-starved *Salmonella* (high *P*_ryhB-2_ activity) resided in macrophages with large or small endosomes (Fig. 2A-D).

**Fig. 2.**
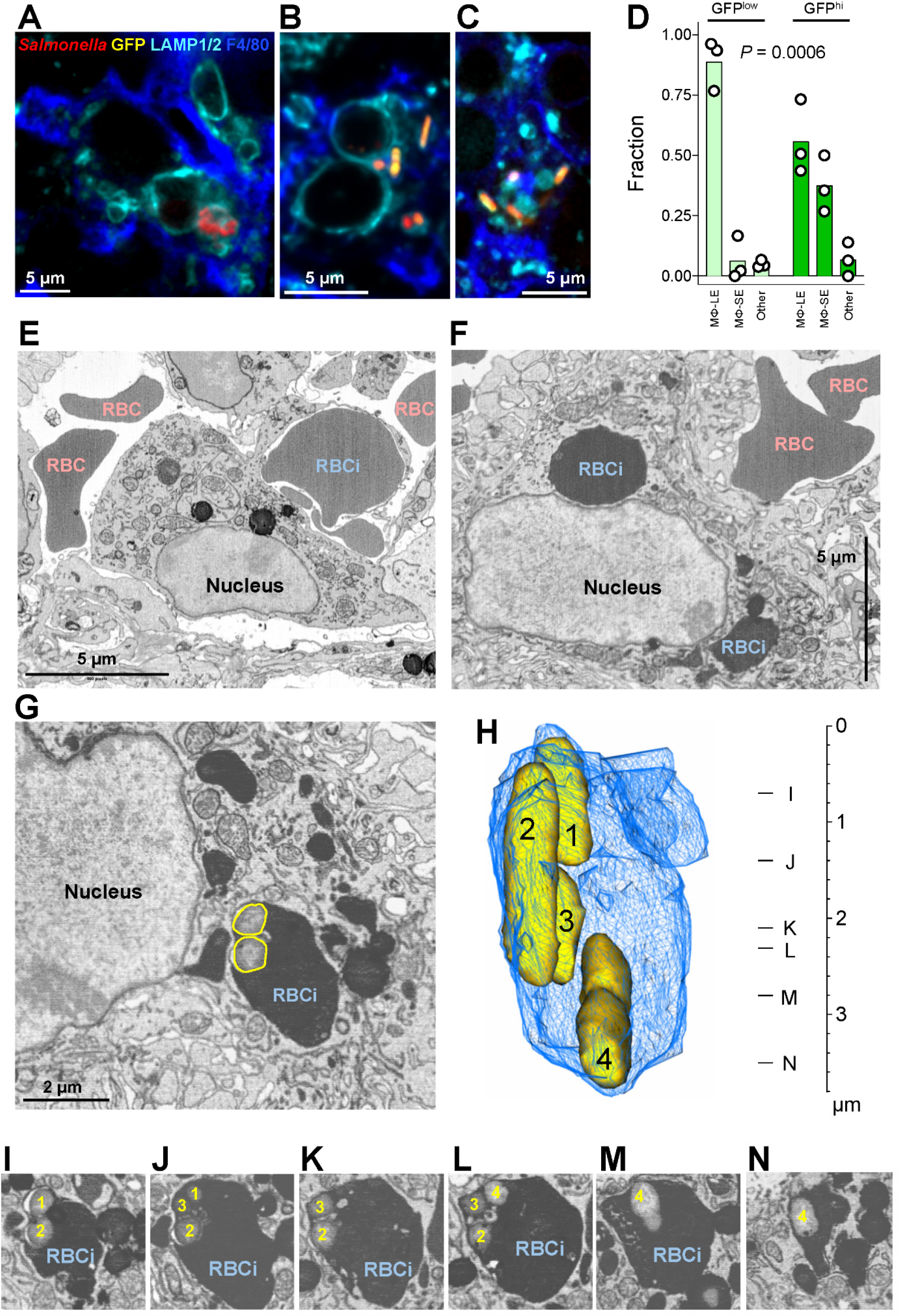
Macrophage erythrophagocytosis delivers iron to some *Salmonella*. **(A)** Confocal fluorescence micrograph of iron-replete *Salmonella* (low *P*_ryhB-2_-*gfp-ova* activity; constitutive mCherry fluorescence) in a F4/80-positive red pulp macrophage that also contains large LAMP1/2-positive endosomes in spleen. **(B)** Iron-starved *Salmonella* (high *P*_ryhB-2_-*gfp-ova* activity; constitutive mCherry fluorescence) in a macrophage that also contains large endosomes. **(C)** Iron-starved *Salmonella* (high *P*_ryhB-2_-*gfp-ova* activity; constitutive mCherry fluorescence) in a macrophage that contains only small endosomes. **(D)** Quantification of iron-replete (GFP^low^) and iron-starved (GFP^high^) *Salmonella* in red pulp macrophages with large endosomes (Mф-LE) or only small endosomes (Mф-SE), or other cells. Each dot represent data from an individual mouse (N: 137, 263, and 300 *Salmonella* cells; two-way ANOVA for test of interaction of GFP level and cell type). **(E)** Serial block-face scanning electron microscopy image of a macrophage phagocytosing a red blood cell (RBCi) in spleen (“erythrophagocytosis”). Free red blood cells (RBC) reside in the interstitial space. **(F)** Macrophage with two ingested red blood cells (RBCi). **(G)** Macrophage with an endosome that contains several *Salmonella* (yellow lines) and material derived from an ingested red blood cell (RBCi). Similar images were obtained for dozens of *Salmonella* in three independently infected mice. **(H)** 3D reconstruction of the *Salmonella*-containing endosome shown in G based on adjacent layers of the 3D tomogram. The numbers indicate the four Salmonella cells in this endosome. The letters indicate the position of layers shown in panels I to N. The scale shows the z-axis which is vertical to the image plane of panels G and I to N.

Serial-blockface scanning electron-microscopy (SBEM) (*35*) revealed that these large macrophage endosomes had homogeneous electron-dense contents indicative of red blood cells (Fig. 2E-G). Erythophagocytosis, the process of capture, digestion and recycling of aged red blood cells, is a key function of red pulp macrophages (*36*) and is enhanced during *Salmonella* infection (*37*).

In SBEM, ex-vivo sorted, live *Salmonella* (*38*) appeared as micrometer-sized rods with an electron-dense cortex and electron-light center (sfig. 5a,b). We detected similar rods in the spleen (sfig. 5c) but some rods had granular electron-dense contents without clear distinction between cortex and center (sfig. 5c) resembling dead bacteria (*39*). This was consistent with detection of ∼40% dead bacteria in spleen using other methods (*38*). Here, we focussed on the microenvironments of live *Salmonella*.

All *Salmonella* were found intracellularly in vacuoles, mostly within macrophages but also in polymorphonuclear neutrophils and other cells (*34*). In macrophages involved in erythrophagocytosis (*40*), some *Salmonella* resided directly in endosomes containing red blood cell-derived contents (Fig. 2G-N; sfig. 5a,b). Since hemoglobin, which constitutes around 97% of the dry mass of red blood cells (*41*), contains four iron atoms per tetramer, these endosomes likely provided abundant iron to *Salmonella*, consistent with the presence of iron-replete *Salmonella* in such cells (Fig. 2A).

Other *Salmonella* in macrophages with erythrophagocytosis were located in electron-light endosomes that showed no connection to red blood cell-derived endosomes (sfig. 5c,d), consistent with the occurrence of iron-starved *Salmonella* in these macrophages despite their high iron content (Fig. 2B). Thus, erythrophagocytosis supplied iron (*42*) to a subset of *Salmonella*, relieving SLC11A1-mediated iron deprivation for that subset.

### Iron-starved *Salmonella* make limited contributions to overall *Salmonella* growth

Iron-starved *Salmonella* might struggle to replicate, limiting their contribution to overall *Salmonella* growth and disease progression. To test this idea, we depleted iron-starved *Salmonella* by inducing expression of the toxin gene *doc* (*43, 44*) during iron starvation using *P*_ryhB-2_. To avoid depleting iron-replete *Salmonella* with baseline *P*_ryhB-2_ activity, we embedded *doc* in a stringent ON/OFF module (*45*) which prevents gene expression unless the promoter activity is high (Fig. 3A; sfig. 6a). Additionaly, constitutive *mCherry* expression allowed us to detect all *Salmonella* cells irrespective of their *P*_ryhB-2_ activity. *Salmonella* carrying an analogous plasmid with an inactive *doc* variant (H66Y D70N A76E) (*43*) and a constitutive *bfp* expression cassette as the control strain (Fig. 3A; sfig.6b). When grown in iron-replete conditions, both strains exhibited similar growth rates, while iron starvation abolished the growth of the strain carrying active *doc* but not the control strain, confirming conditional depletion (sfig. 6c).

**Fig. 3.**
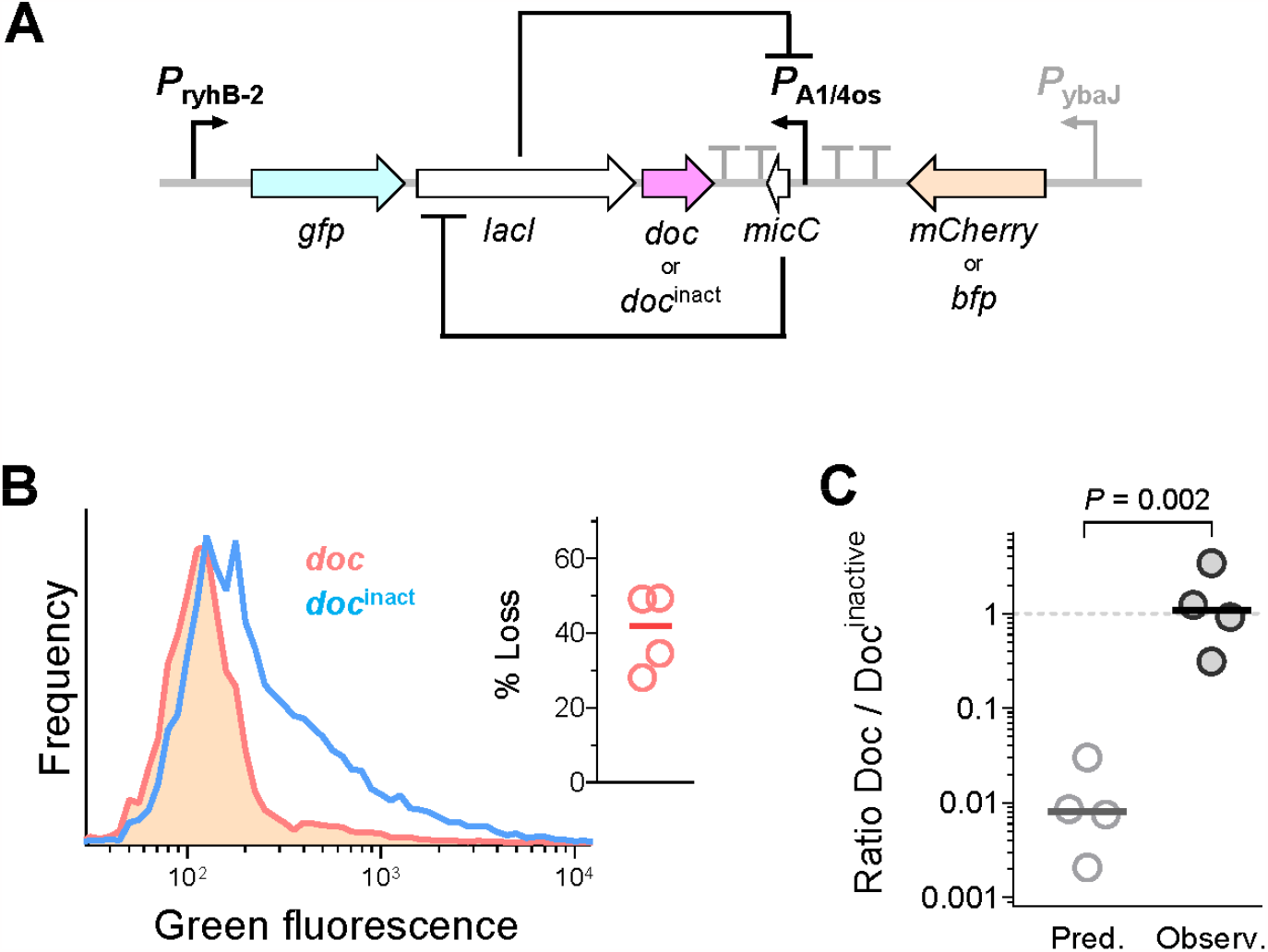
Limited contribution of the iron-starved subset to overall *Salmonella* growth. **(A)** Construct for arresting growth of iron-starved *Salmonella. P*_ryhB-2_ drives expression of *gfp-ova* encoding an unstable variant of the green fluorescent protein. Only high *P*_ryhB-2_ activities can overcome the silencing of the toxin gene *doc* by the stringent ON/OFF module consisting of two inhibitory feedback loops mediated by *lacI* encoding the *lac* repressor which inhibits the *P*_A1/4os_ promoter which drives expression of the small non-coding rna Mic, which represses *lacI* translation.Iron-regulated *doc* expression thus causes growth inhibition exclusively in cells with iron starvation inducing high *P*_ryhB-2_ activity. For detection of all Salmonella regardless of *P*_ryhB-2_ and Doc levels, *mCherry* is expressed from the constitutive *P*_ybaJ_ promoter. As a control, we used inactive *doc* H66Y D70N A76E coupled with constitutive *bfp* expression (instead of *mCherry)*. **(B)** Green fluorescence of the test strain with conditional expression of active *doc* (red) and the control strain with inactive *doc* (blue) in spleen of C57Bl/6 *SLC11A1*^r/r^ mice. Pooled data for four independently mice are shown. The inset shows the loss of bright cells in the test strain for individual mice. **(C)** Predicted and observed competitive growth of test and control strains in mixed infections. Based on ∼40% loss and ten divisions, the test strain could grow poorly. However, the experimental data do not show and growth difference between test and control strains (paired t-test on log-transformed data).

We infected C57BL/6 *SLC11A1*^r/r^ mice with a mixture of both strains and observed colonization of the spleen by both strains (sfig. 6d). The blue-fluorescent control *Salmonella* had normal broad *P*_ryhB-2_ activities (Fig. 3B). The red *Salmonella* exhibited mostly low *P*_ryhB-2_ activity while the subset of ∼40% with high activity was lacking, confirming conditional depletion in vivo (Fig. 3B).

If iron-replete and iron-starved *Salmonella* had similar fitness, the depletion of 40% iron-starved cells would reduce the overall *Salmonella* load ∼150-fold: instead of 2 daughter cells per division, an average of 1.2 cells would survive. After ∼10 net divisions during six days of infection (*18*), the strain carrying active *doc* would increase only by a factor of 1.2^10^ = 6.2-fold, whereas the control strain would increase by a factor of 2^10^ = 1024-fold (Fig. 3C). Contrary to these predictions, active-*doc* and control *Salmonella* reached similar loads after six days of infection (Fig. 3C), indicating that depletion of the iron-starved subset had limited overall impact. This suggests that iron-replete *Salmonella* generate the majority of offspring with only small contributions form iron-starved *Salmonella*.

### Iron starvation controls replication of a minor *Salmonella* subset

To investigate if this limited contribution of iron-starved *Salmonella* was due to poor replication, we determined the division rates of *Salmonella* with different *P*_ryhB-2_ activities using the single-cell growth reporter TIMER^bac^ (*18, 46*) (sfig. 8). In mice with defective iron transporter SLC11A1, *Salmonella* exhibited baseline *P*_ryhB-2_ activity and a median division rate of 0.13 ± 0.01 h^-1^ (*18*) (Fig. 1C; Fig. 4A,D,E).

**Fig. 4.**
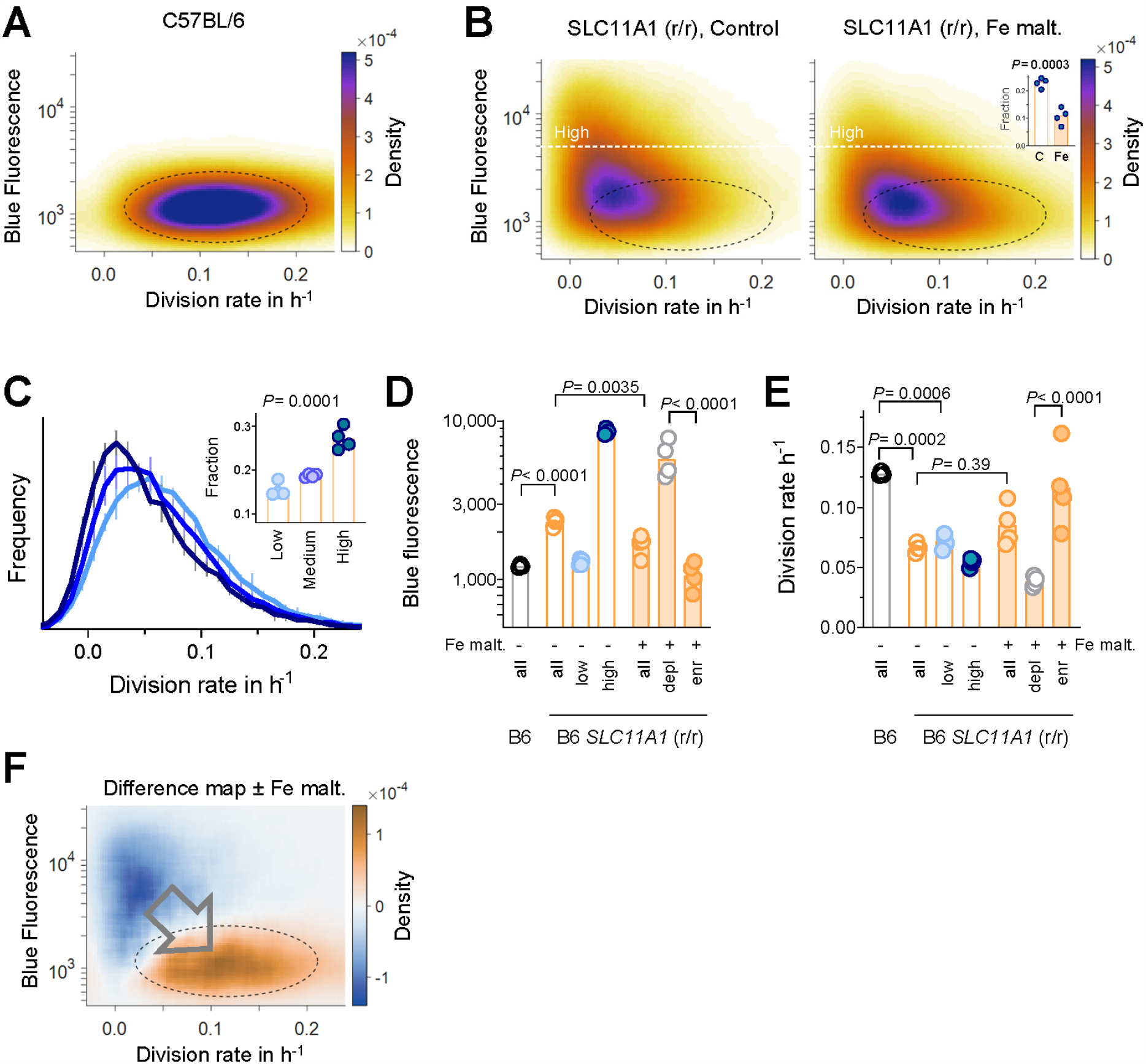
Growth-limiting iron-starvation in a minor *Salmonella* subset. **(A)** Relationship between division rate and *P*_ryhB-2_*-bfp-ova* activity (as measured by blue fluorescence) of single *Salmonella* cells in C57BL/6 mice. Pooled data of three mice are shown (the blue fluorescence data are also shown in Fig. 1C,D and 4D). The dashed black ellipse contains 70% of all *Salmonella* cells. Median division rates for individual mice are shown in panel E. **(B)** Relationship between *Salmonella* division rate and *P*_ryhB-2_*-bfp-ova* activity in C57BL/6 *SLC11A1*^r/r^ mice without (Control) or with excess iron supplementation (Fe malt.). Pooled data for four mice in each condition are shown. The black ellipse shows the core population of *Salmonella* in C57BL/6 mice (panel A). The white dashed line represents our threshold (5,000) of *Salmonella* with high blue fluorescence. The inset shows the fraction of cells with high blue fluorescence in control (C) and iron-supplemented (Fe) mice (paired t-test). Median blue fluorescence and division rates are shown in panels 4D,E. **(C)** Division rates of *Salmonella* with low *P*_ryhB-2_*-bfp-ova* activity (blue fluorescence<2,000), medium activity (blue fluorescence between 2,000 and 5,000), or high activity (blue fluorescence >5,000) in C57BL/6 *SLC11A1*^r/r^ mice with normal iron nutrition. Pooled data for 4 mice are shown. The inset shows the fraction of growth-arrested *Salmonella* (apparent division rate <0.02 h^-1^) for individual mice (one-way ANOVA with test for linear trend). **(D)** Blue fluorescence of Salmonella in C57BL/6 mice (B6) or C57BL/6 *SLC11A1*^r/r^ mice without/with excess iron supplementation (Fe malt.). Medians of individual mice and their entire *Salmonella* population (all) and subsets with low or high blue fluorescence are shown. For mice with excess iron supplementation, the disappearing subsets (depl) and the merging subsets (enr) are shown. **(E)** Division rates for the same *Salmonella* populations as shown in D. **(F)** Difference map of division rate and *P*_ryhB-2_*-bfp-ova* activity for mice with excess iron supplementation (Fe malt.) compared to control mice. Pooled data for four pairs of mice are shown. Median data for individual mice are shown in panels D and E.

In mice with restored SLC11A+, *Salmonella* showed heterogeneous *P*_ryhB-2_ activity (Fig. 1C, Fig. 4B,D,E). Some *Salmonella* had low activities, indicating that they had easy access to iron similar to that in C57BL/6 mice, yet they still divided slowly (0.07 ± 0.01 h^-1^). This was consistent with the finding that, in the presence of SLC11A1, magnesium rather than iron is the limiting factor for most *Salmonella* (*18*) (Fig. 4C). On the other hand, severely iron-starved *Salmonella* with high *P*_ryhB-2_ activity divided even more slowly (0.055 ± 0.01 h^-1^) and ∼a quarter of them showed growth arrest (division rate< 0.02 h^-1^, equivalent to generation times >50 h) (Fig. 4C). These data might still underestimate the fraction of growth-arrested cells because TIMER^bac^ only responds after ∼one generation time, resulting in delayed detection of recently decelerating bacteria (*46*). Thus, iron-starved *Salmonella* replicated less than iron-replete *Salmonella*, consistent with their limited contribution to overall *Salmonella* growth.

This impaired replication could be caused directly by poor access to iron, or other factors that coincide with low iron availability. To test the role of iron limitation, we administered iron isomaltoside, a clinically approved complex for slow release of bioavailable iron directly to macrophages (*47*). This iron supplementation reduced overall *P*_ryhB-2_ activities and decreased the fraction of severely iron-starved *Salmonella* ∼2-fold (Fig. 4B,D,E), indicating effective iron delivery to starving *Salmonella*. However, the overall impact on *Salmonella* division rates was not statistically significant (Fig. 4A-C). Spleens of supplemented mice contained only 3.3 ± 1.1-fold more *Salmonella* than control mice at day six post-infection *(P =* 0.0017; one-sample t-test on log-transformed data). This difference was relatively small compared to the ∼1,000-fold increase in *Salmonella* loads during the same infection period. Thus, the overall impact of excess iron was minimal.

However, single-cell analysis revealed heterogeneous *Salmonella* responses (Fig. 4F). While most *Salmonella* were unaffected, a subset of ∼20 % otherwise iron-starved *Salmonella* showed a ∼5-fold reduction in *P*_ryhB-2_ activity and a ∼3-fold increased division rate, resembling the properties of *Salmonella* in C57BL/6 mice lacking functional SLC11A1 (Fig. 4A,D-F). Thus, SLC11A1 indeed limited a minor *Salmonella* subset by iron deprivation, which could be relieved by an excess of iron. However, this iron-responsive minor subset was overshadowed by the majority of *Salmonella* with easy access to iron by limiting magnesium starvation (*18*).

## Conclusion

The impact of iron on *Salmonella* infections remains poorly understood despite extensive research. Our findings demonstrate that commonly studied mouse strains, which exhibit disrupted iron homeostasis, do not impose significant iron limitations on *Salmonella*. Hence, iron perturbations primarily affect host immune functions rather than *Salmonella* nutrition as widely assumed. Under more physiological conditions, the host iron transporter SLC11A1 induces growth-limiting iron starvation in a minor *Salmonella* subset, thereby restricting its contribution to overall *Salmonella* growth. However, this subset is masked by the majority of *Salmonella* which access iron-rich endosomes in macrophages that degrade red blood cells, effectively bypassing the iron limitation imposed by SLC11A1. Nonetheless, SLC11A1 still hinders replication of these *Salmonella* through magnesium deprivation (*18*). Although SLC11A1 also reduces *Salmonella*’s access to zinc, this does not significantly impact replication (*18*). Thus, a heterogeneous, magnesium-iron axis of nutritional immunity controls *Salmonella* infection. Our results highlight the importance of single-cell analysis in physiologically relevant animal models for unraveling complex mechanisms of infection control.

## Acknowledgements

We would like to thank Jasmin Künneke for help with analyzing imaging data, Janine Bögli (Biozentrum FACS Core Facility) for sorting, Carola Alampi and Cinzia Tiberi (Biozentrum BioEM lab) for scanning electron micrographs of *Salmonella*, Christel Genoud (Friedrich-Miescher-Institute) for help with serial block-face scanning electron microscopy, and Alexander Hoffmann (Innsbruck) for help with iron maltoside;

## Author contributions

Outlined the study: DB, BR, Performed experiments: BR, OC, CB, BC, MAV, JL, Analyzed data: BR, OC, BC, CB, MAV, DB, Interpreted data: BR, OC, CB, DB, Wrote manuscript: DB, BR;

## Data and materials availability

All data is available in the manuscript or the supplementary materials.

Correspondence and requests for materials should be addressed to D.B.

## Supplementary Materials

Materials and Methods

Figs. S1-S8

Supplemental References (1-14)

