## Supplemental data for "Heterogeneous dual-metal control of *Salmonella* infection"

**This PDF file includes:**

Materials and Methods

Figs. S1 to S8

Supplementary References

**Materials and Methods**

### Bacterial strains, plasmids and growth media

*Salmonella* strains used in this study were based on SME51, a prototrophic *hisG*^Leu69^ derivative of *Salmonella enterica* serovar Typhimurium SL1344 (*1-3*).

Reporter strains carried low-copy episomal pSC101 derivatives (sfig. 4a, 7a,b, 8) with

constitutive expression of *mCherry*, *mtagbfp2* (*4*), or *timer*^bac^ (*3*) from the *P*_ybaJ_ promoter (*5*), and fusions of promoters *P*_mgtCQBRUcigR_ (chromosomal coordinates in SL1344 GenBank: FQ312003.1: 3,987,050 - 3,986,453) or *P*_ryhB-2_ (1,309,793 – 1,310,035) driving expression of *gfp-ova* coding for a degradable variant of the green fluorescent protein GFP.mut2 (*6, 7*) or expression of *mtagbfp2-ova*.

Bacteria were grown in Lennox lysogeny broth (LB) or in MES-MM (100 mM MES-KOH pH 5.5, 0.4% glycerol, 15 mM NH4Cl, 1.5 mM K2SO4, 3 mM KH2PO4) containing 90 mg/L streptomycin and 50 mg/L kanamycin. For metal depletion, MES-MM was treated with Chelex100 for 4h. In-vitro growth in 96-well plates (Greiner, PS flat-bottomed microplate) at 37°C with shaking was monitored by OD_600_ measurements at 15 min intervals in a Epoch2 microplate reader (BioTek).

### Mouse Infection

All animal experiments were approved (license 2239, Kantonales Veterinäramt Basel) and performed according to local guidelines (Tierschutz-Verordnung, Basel) and the Swiss animal protection law (Tierschutz-Gesetz). 10 to 15 week-old BALB/c, C57BL/6, or C57BL/6 *SLC11a1*^r,r^ (*8*) mice were infected by tail-vein injection of *Salmonella* strain mixtures containing 2,000 to 5,000 CFU in 100 µl PBS. The inoculum size was determined by plating for each infection. We used both female and male mice. Four (BALB/c, C57BL/6) or six days (C57BL/6 *SLC11a1*^r,r^) post-infection, the mice were euthanized with CO_2_ and the spleen was prepared. *Salmonella* load was determined by plating on lysogeny broth agar. The “competitive index” (CI) was determined by dividing the output ratio by the inoculum ratio. Some C57BL/6 SLC11a1^r,r^ mice received 1 mg of iron maltoside (Monofer, Pharmacosmos) dissolved in 100 μl 0.9 % sodium chloride solution by tail-vein injection four days prior to the infection. This dose corresponds to ~50 mg/kg compared to a single dose of 20 mg/kg recommended for treating human patients with iron deficiency. Control mice received only vehicle.

We estimated sample size by a sequential statistical design. We first infected two to three mice based on effect sizes and variation observed in our previous studies (*9*), and used the results to estimate group sizes for obtaining statistical significance with sufficient power. The experiments were neither randomize nor blind. However, flow cytometry analysis was carried out using an automated unbiased approach (see Flow Cytometry section).

### Flow cytometry

The spleen was homogenized in ice-cold phosphate-buffered saline containing 0.2% Triton X-100. All samples were kept on ice until and during analysis. Large host cell fragments were removed by repeated centrifugations at 100xg for 10 min at 4°C. Relevant spectral parameters were recorded in a FACS Fortessa II (Becton Dickinson), using thresholds on SSC to exclude electronic noise. We used the following channels: mTagBFP2, excitation 405 nm, emission 460-480 nm (“blue”); GFP and green TIMER^bac^ component, excitation 488 nm, emission 502-525 nm (“green”); mCherry and red TIMER^bac^ component, excitation 561 nm, emission 595-664 nm (“red”); orange autofluorescence, excitation 445 nm, emission 573-613 nm. All parameters were measured as “height”. Fluorescent *Salmonella* cells were purified from spleen homogenates using a FACSAria Fusion cell sorter (BD Biosciences) using the same channels as above. Data were processed with FlowJo and MATLAB. TIMER^bac^ fluorescence log color-ratios were converted into *Salmonella* division rates based on previously established calibration data (*3*).

### Proteomics

Sorted *Salmonella* cells (1 to 6.5 Mio) were resuspended in lysis buffer (5% sodium dodecyl sulfate, 10mM tris(2-carboxyethyl) phosphine, 100 mM triethyloammonium bicarbonate) and incubated at 95°C for 10 min. Samples were then sonicated with 20 cycles On/Off cycles (30 s, 30 s) using a Bioruptor

(Diagenode). The protein content was determined by tryptophan-based fluorescence assay (Infinite M Plex, Tecan). Samples were alkylated by addition of 20 mM iodoacetamide and incubation at 25°C for

30 min with gentle shaking. Material containing 20 μg protein was processed using S-Trap (ProtiFi). Peptide concentration was determined using a UV-based assay (Infinite M Nano, Tecan). Peptides were separated on a Dionex UltiMate 3000 system (ThermoFisher Scientific) coupled online to an Orbitrap Exploris 480 mass spectrometer (ThermoFisher Scientific) using ID capillary columns (20 cm long, diameter 75 μm) packed in-house with 1.9 μm Reprosil-Pur C18 beads (Dr. Maisch, Ammerbuch, Germany). The column temperature was maintained at 50 °C using an integrated column oven interfaced online with the mass spectrometer. Formic acid (FA) 0.1% was used to buffer the pH in the two running buffers used. The total gradient time was 60 min and went from 2% to 12% acetonitrile (ACN) in 5 min, followed by 45 min to 35%, and 10 min at 50%. This was followed by a washout by 95% ACN, which was kept for 20 min, followed by re-equilibration with 0.1% FA buffer. Flow rate was kept at 300 nL/min. Spray voltage was set to 2,500 V, funnel RF level at 40, and heated capillary at 275 °C. For data-independent acquisition full MS resolution was set to 120,000 and full MS normalized AGC target was 300% with an IT of 45 ms. Mass range was set to 350–1400. Normalized AGC target value for fragment spectra was set at 1000%. A total of 63 windows of 9 Da were used with an overlap of 1 Da. Resolution was set to 15,000 and IT to 22 ms. Normalized CE was set at 28%. All data were acquired in centroid mode using positive polarity and advanced peak determination was set to on.

For data processing and protein identification, raw data were imported into SpectroNaut (16.1.220730.53000, Biognosys) and analyzed with directDIA. Searches were carried out against a fasta file including the proteomes of *Salmonella enterica* serovar Typhimurium SL1344 and *Mus musculus*. Cysteine carbamidomethylation was set as fixed modification. Methionine oxidation, methionine excision at the N terminus were selected as variable modifications. Results were filtered for a 1% false discovery rate (FDR) on spectrum, peptide, and protein levels. Mass spectrometry data have been deposited to the ProteomeXchange Consortium via the PRIDE (30395289) partner repository with the data set identifier PXDXXXXXX. Proteomics data analysis was performed with Perseus (1.6.14.0). Protein abundance was calculated using protein group quantities (PG quantity) based on the “total protein approach” concept (*10*). Statistical analysis was based on data permutations and significanceanalysis-of-microarrays statistics (44) using a s0 value of 0.1**.**

### Immunohistochemistry and confocal microscopy

Infected mice were anesthetized with 100 mg/kg ketamine and 16 mg/kg xylazine. Mice were transcardially perfused with 15 ml ice-cold PB buffer (0.1 M phosphate buffer pH 7.2 without NaCl) containing 10 U/ml heparin, followed by 35 ml 4% paraformaldehyde (PFA) in PB with heparin. Perfusion-fixed spleen was prepared, cut into four pieces, and post-fixed in 4% PFA in PB overnight.

Samples were washed thrice in PB for 15 min at 4°C and embedded in 4% ROTI®Garose with low melting and gelling temperature (Roth) in PB buffer. Spleen sections with 50 μm thickness were cut using a Compresstome (Precisionary). The sections were put in ice-cold cryoprotectant (300 g sucrose, 10 g polyvinyl-pyrrolidone 40, 400 ml ethylene glycol, 400 ml PB, after dissolving add PB to 1 l) (*11*). One hour after sections had sunken to the bottom of the tube, the tube was placed for at least seven days in a freezer at -20°C.

Sections were warmed to room temperature and washed for 5 min with TBS (20 mM Tris base, 150 mM NaCl, adjusted to pH 7.4 with HCl)). For antigen retrieval, sections were incubated in prewarmed 10 mM sodium citrate pH 8.5 at 37°C for 45 min. Sections were washed with TBST (0.1% Triton X-100 in TBS), blocked for 1 h with 10 mg/mL BSA-1 % mouse serum in TBST, and washed again in TBST. Sections were incubated with 10 µg/mL biotinylated F4/80 (clone Cl:A3-1, AbD Serotec MCA497B), 4 µg/mL anti-CD107a (LAMP-1)-BV421 (clone: 1D4B, Biolegend 121617), and 4 µg/mL anti-CD107a (LAMP-1)-BV421 (clone: ABL-93, BD 564249) at 4°C overnight, washed in TBST, and stained with 10 µg/mL Streptavidin- Alexa 647 (ThermoFischer S21374) for 1 h at room temperature.

Sections were washed in TBS and mounted in mounting medium (DAKO). Imaging was done with a SpinSR spinning-disk confocal super-resolution microscope (Olympus) equipped with a GFP narrow band emission filter (500-520 nm) using the 60x objective with additional 3.2x magnification and Super Resolution mode (back-projected pinhole radius 130 nm, distance 2.53 μm). Images were deconvolved with Huygens Professional with the “standard” profile and default settings from the *MOSAIC* file.

Images were analyzed with ImageJ 1.53q (*12*).

### Serial block-face scanning electron microscopy

Samples were prepared according to the protocol developed by the National Center for Microscopy and Imaging Research, University of California, San Diego (https://ncmir.ucsd.edu/sbem-protocol ). In brief, mice were perfusion-fixed with 2.5% glutaraldehyde / 2% paraformaldehyde. Spleen pieces were stained with osmium tetroxide, uranyl acetate, and lead aspartate. After dehydration, tissues were embedded in Durcupan resin. Resin blocks were imaged using a serial block-face scanning electron microscope (Merlin 3View; Zeiss) with an imaging area of 830 x 830 μm^2^, a pixel size of 10 x 10 nm^2^ and slices of 70 nm. Image stacks of 300 to 700 slices were aligned with ImageJ 1.53q (*12*) using the plugin MultiStackReg (*13*). Image analysis and 3D reconstruction was done with 3dmod 4.12.32 (*14*).

### Statistics

Statistical tests were performed with GraphPad Prism 9.3.1 as indicated in the figure legends.

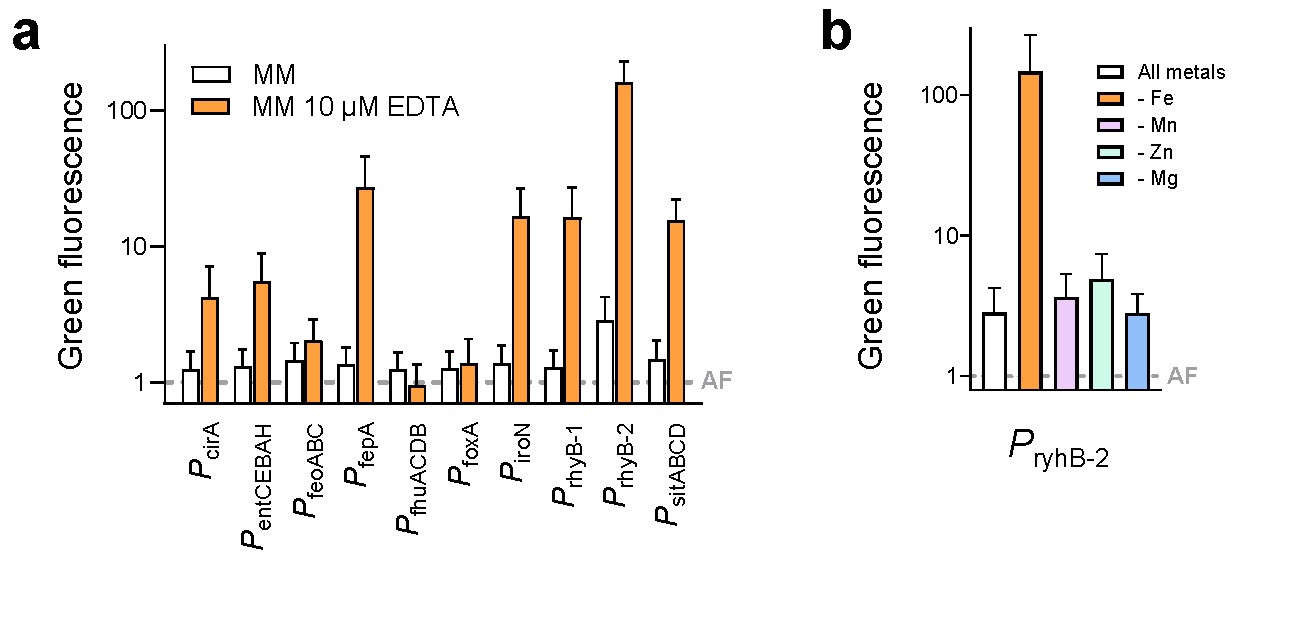

**fig. S1. Iron-responsive *Salmonella* reporter strains**

(a) Green fluorescence levels of *Salmonella* carrying FUR-dependent promoter fusions to *gfp-ova* encoding an unstable GFP variant (*7*) in minimal medium

(MM) or the same medium with 10 µM EDTA. The dashed line represents the *Salmonella* autofluorescence background (AF). Means and SDs of three independent experiments are shown.

(b) *P*_ryhB-2_-*gfp-ova* activity in divalent cation-depleted minimal medium with addback of metals except iron, manganese, zinc, or magnesium. Means and SDs of three independent experiments are shown (AF, autofluorescence).

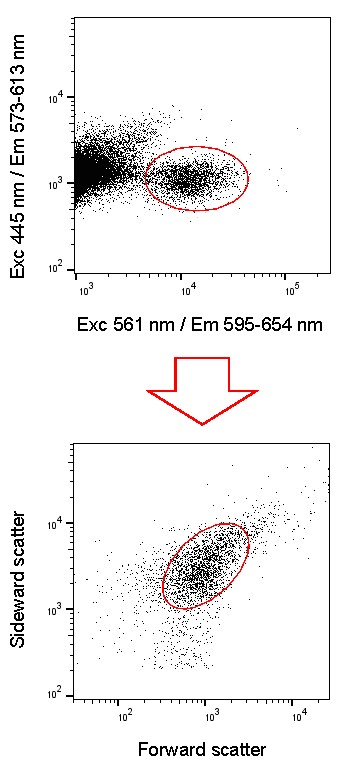

**fig. S2. Gating strategy for identifying *Salmonella* in spleen homogenates.**

Flow cytometry data are shown with each dot representing a particle. All light parameters were measured as signal “height” values (Exc, excitation wavelength; Em, emission range). All *Salmonella* expressed *mCherry* enabling their detection based on green-excited red mCherry fluorescence vs. violet-excited red autofluorescence (upper panel). Additional background was removed based on forward and sideward scatter (lower panel).

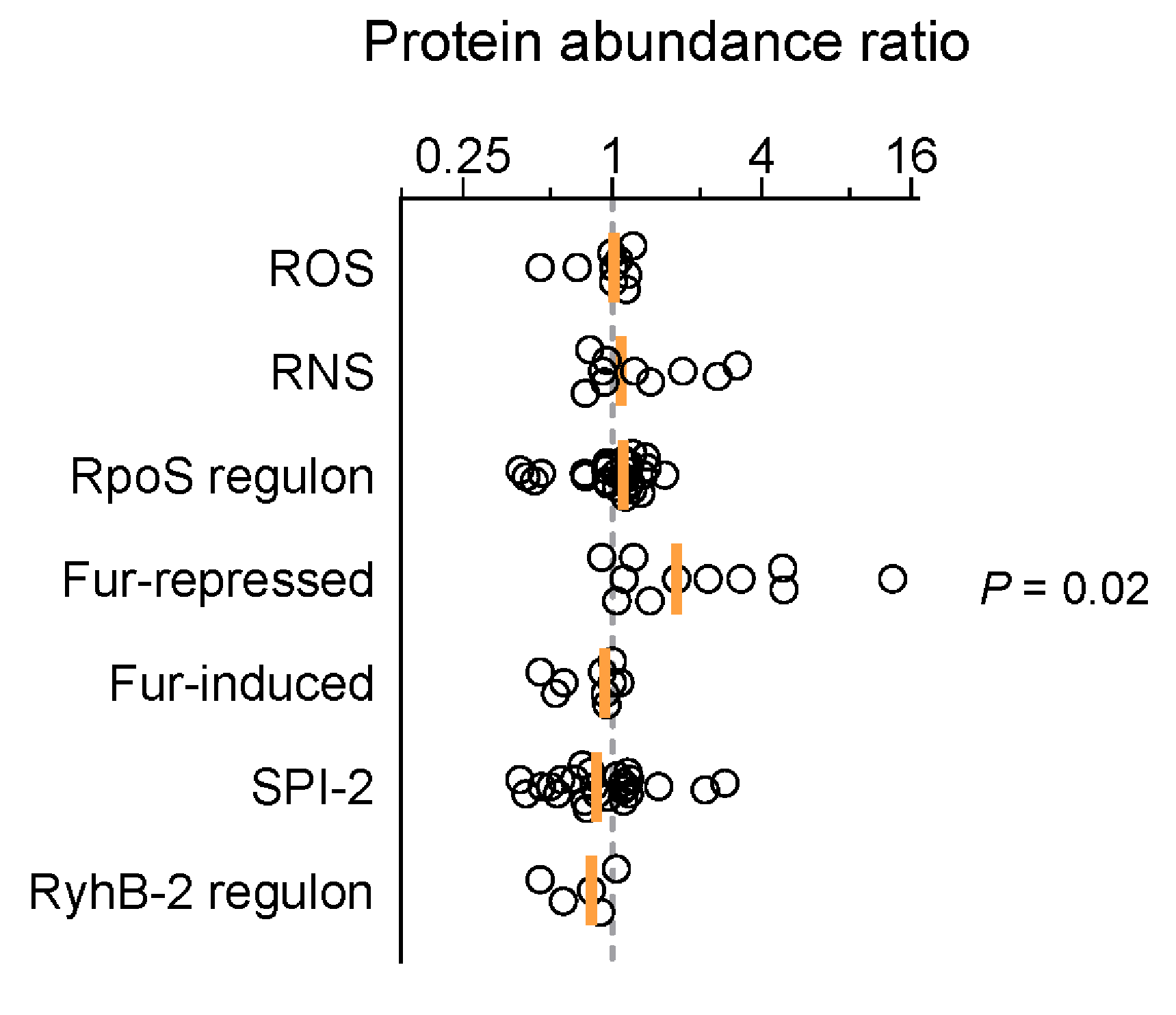

**fig. S3. Protein abundance in *Salmonella* subsets with low or high *P*_ryhB-2_ activity**.

Abundance ratios for proteins associated with oxidative (ROS) or nitrosative (RNS) stress, the starvation-activated sigma factor σ^S^ (RpoS regulon), the iron uptake regulator Fur, virulence genes associated with the type 3 secretion system encoded on *Salmonella* pathogenicity island 2 (SPI-2), and proteins that can be regulated by the small non-coding RNA RyhB-2 (onesample Wilcoxon signed rank test with adjusted *P*-values for multiple testing according to Holm-Šídák).

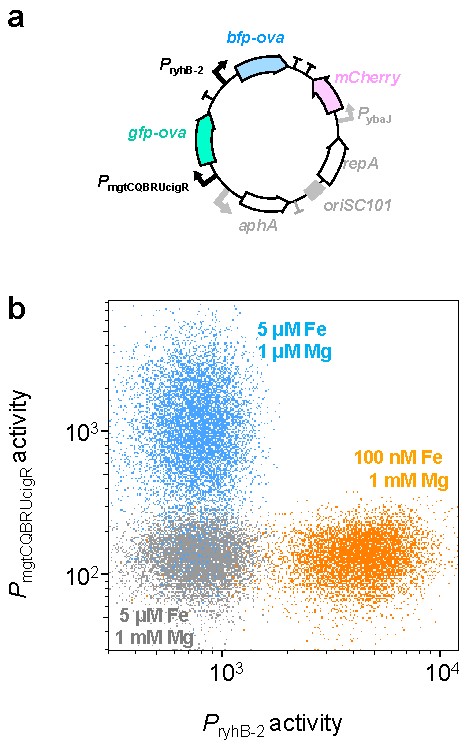

**fig. S4. Dual reporter construct for iron and magnesium access**.

(a) Plasmid map of the three-color construct (*aphA* encodes aminoglycoside phosphotransferase A which confers resistance to kanamycin; *repA* codes for replicase recognizing the origin of replication, *oriSC101*; T, terminator). *mCherry* is expressed from the constitutive promoter *P*_ybaJ_ XXX to identify all *Salmonella* cells against host background (sfig. S1) regardless of GFP and BFP levels.
(b) In-vitro characterization of *Salmonella* carrying the dual reporter plasmid at different iron and magnesium concentrations. An independent experiment showed similar results.

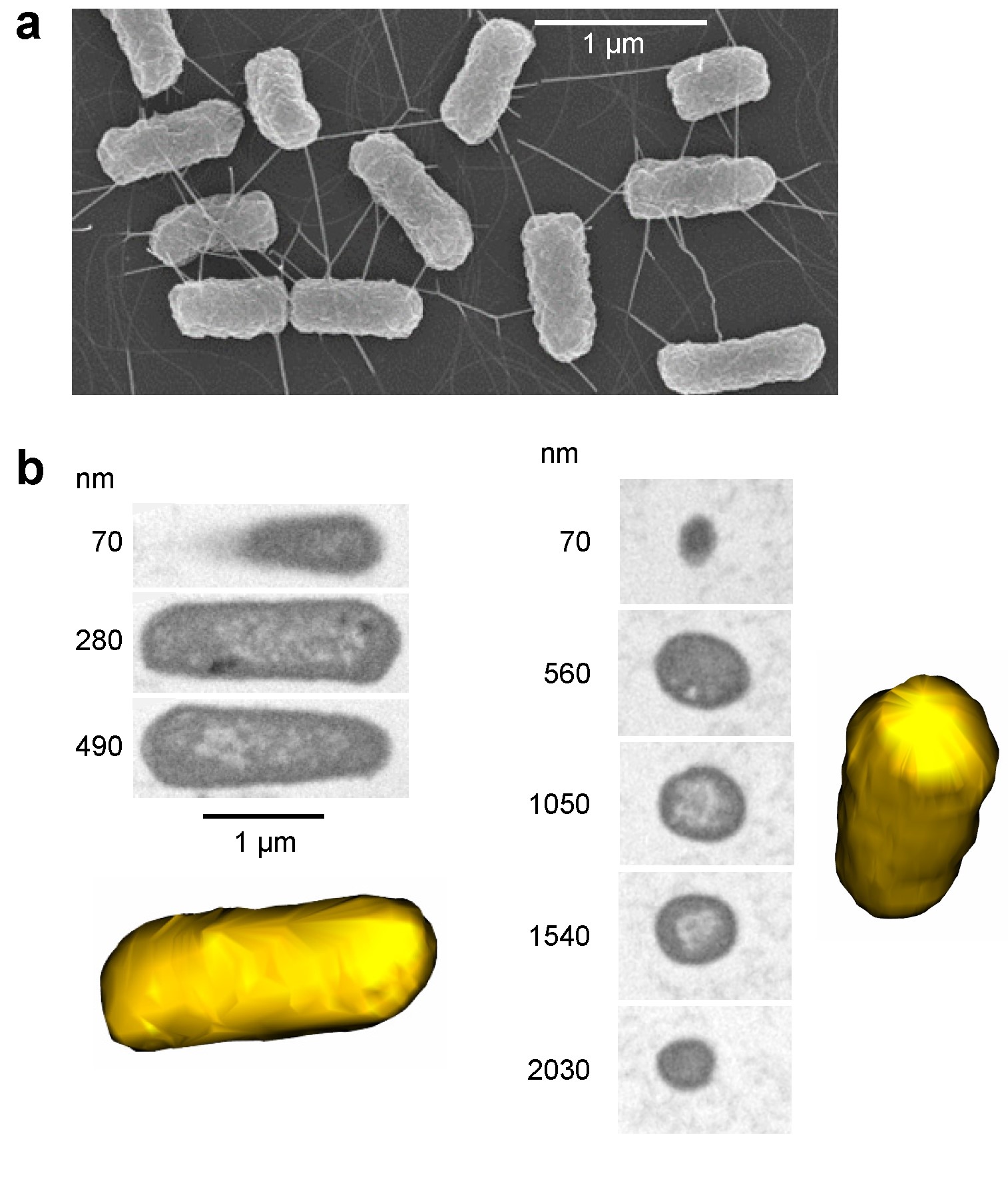

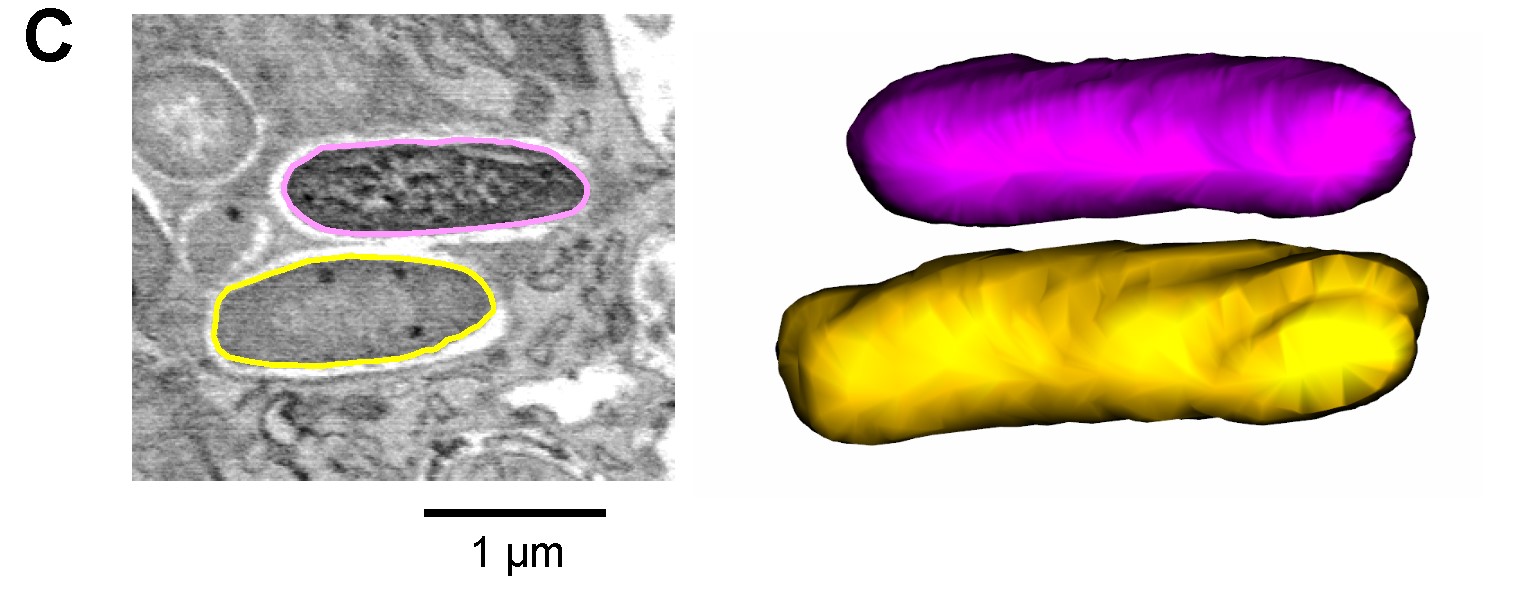

**fig. S5. Identifying of *Salmonella* with serial block-face scanning electron microscopy**. **(a)**	Conventional scanning electron micrograph of a *Salmonella* in-vitro culture showing the typical rod shape of these bacteria.

(b) Serial block-face scanning electron microscopy images of *Salmonella* sorted from infected mouse spleen. Individual layers at different depth and the corresponding 3D reconstructions are shown for two representative *Salmonella* cells.
(c) Serial block-face scanning electron microscopy image of two *Salmonella* cells in mouse spleen. One cell (yellow line) has an electron-dense cortex and an electronlight core similar to the live ex-vivo sorted *Salmonella* cells shown in b. The other cell (magenta line) has granular electron-dense contents with no detectable cortexcore structure resembling images of killed bacteria.

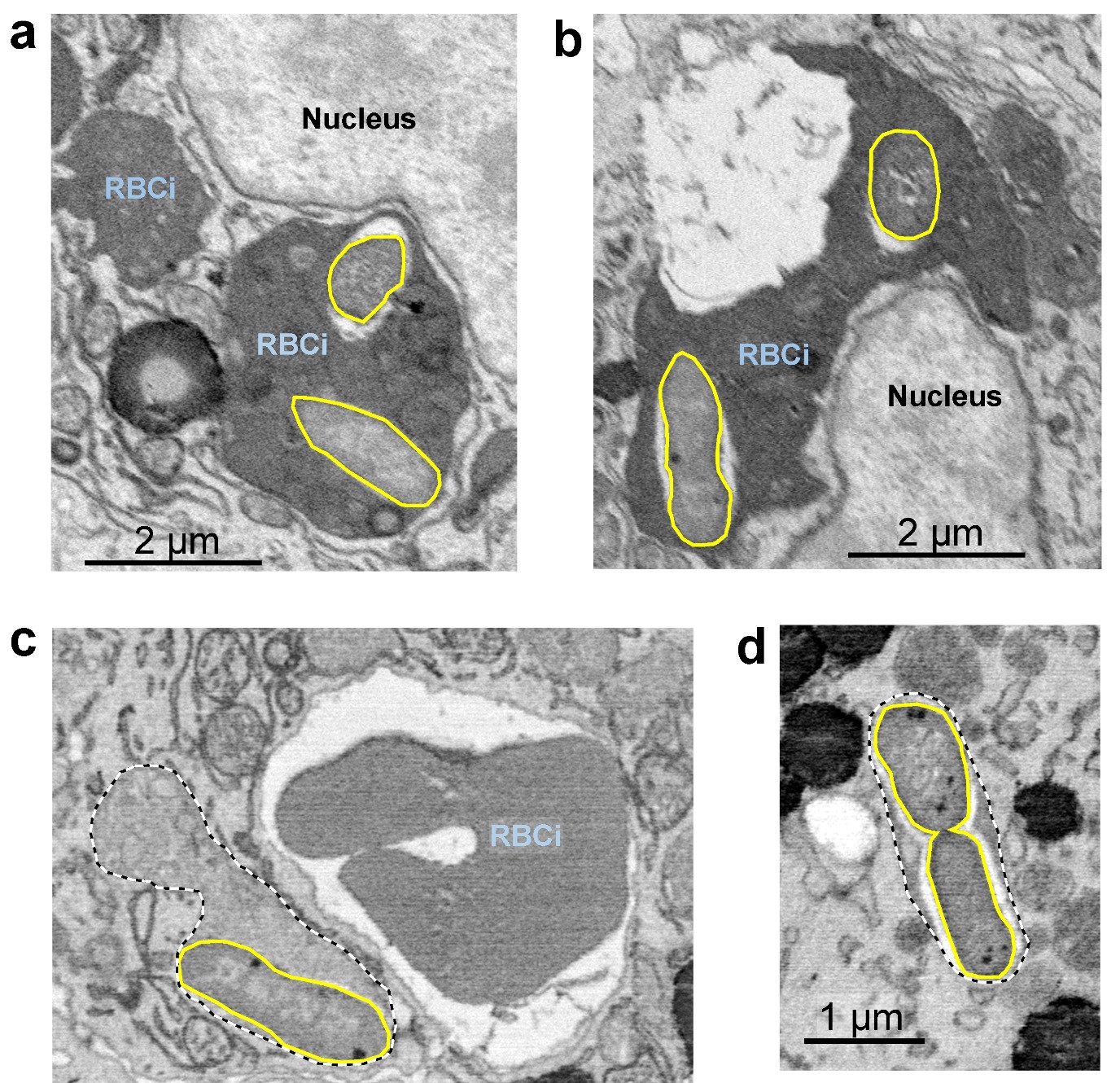

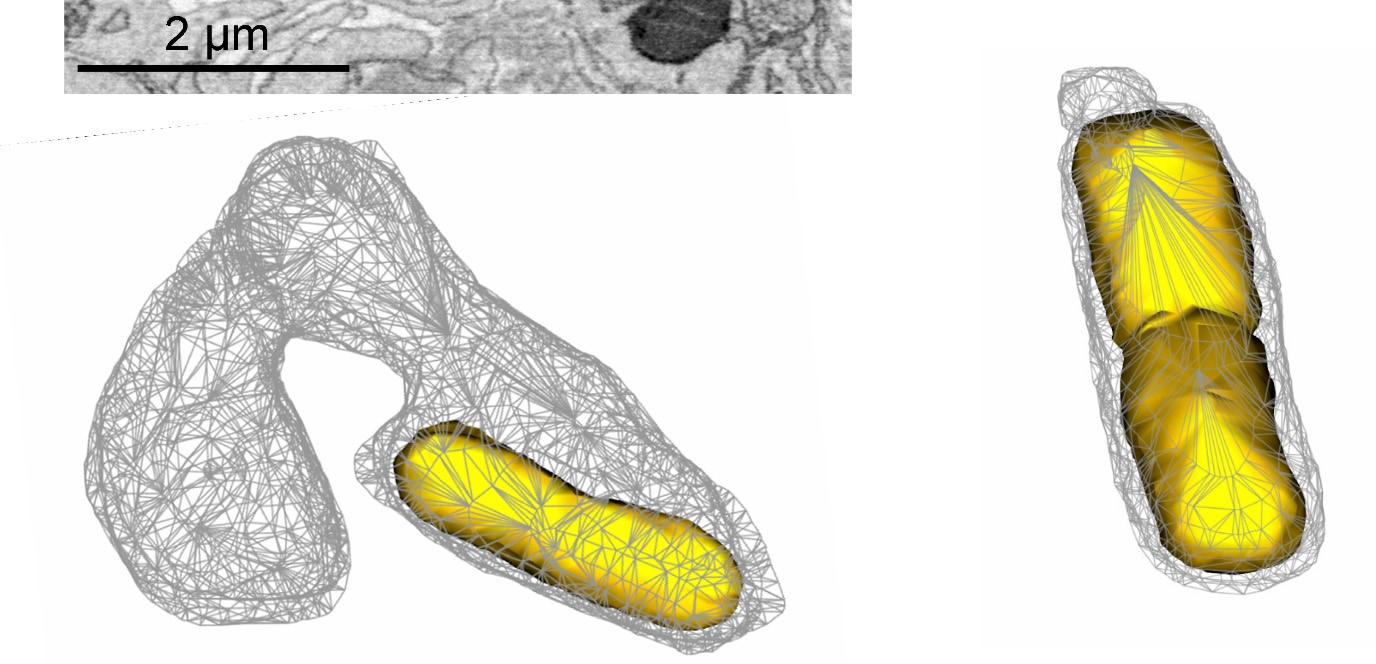

**fig. s6. Serial block-face scanning electron microscopy of *Salmonella* in spleen**. **(a)**	Endosome that contains *Salmonella* (yellow lines) and material derived from an ingested red blood cell (RBCi).

(b) Another endosome in another macrophage that contains *Salmonella* (yellow lines) and material derived from an ingested red blood cell (RBCi).
(c) *Salmonella*-containing endosome in a macrophage that ingested red blood cells

(RBCi). The 3D reconstruction from adjacent layers (below) reveals that the *Salmonella*-endosome has no connection to the adjacent red bllod cell-containing endosome.

(d) *Salmonella*-containing endosome with small volume and little additional material.

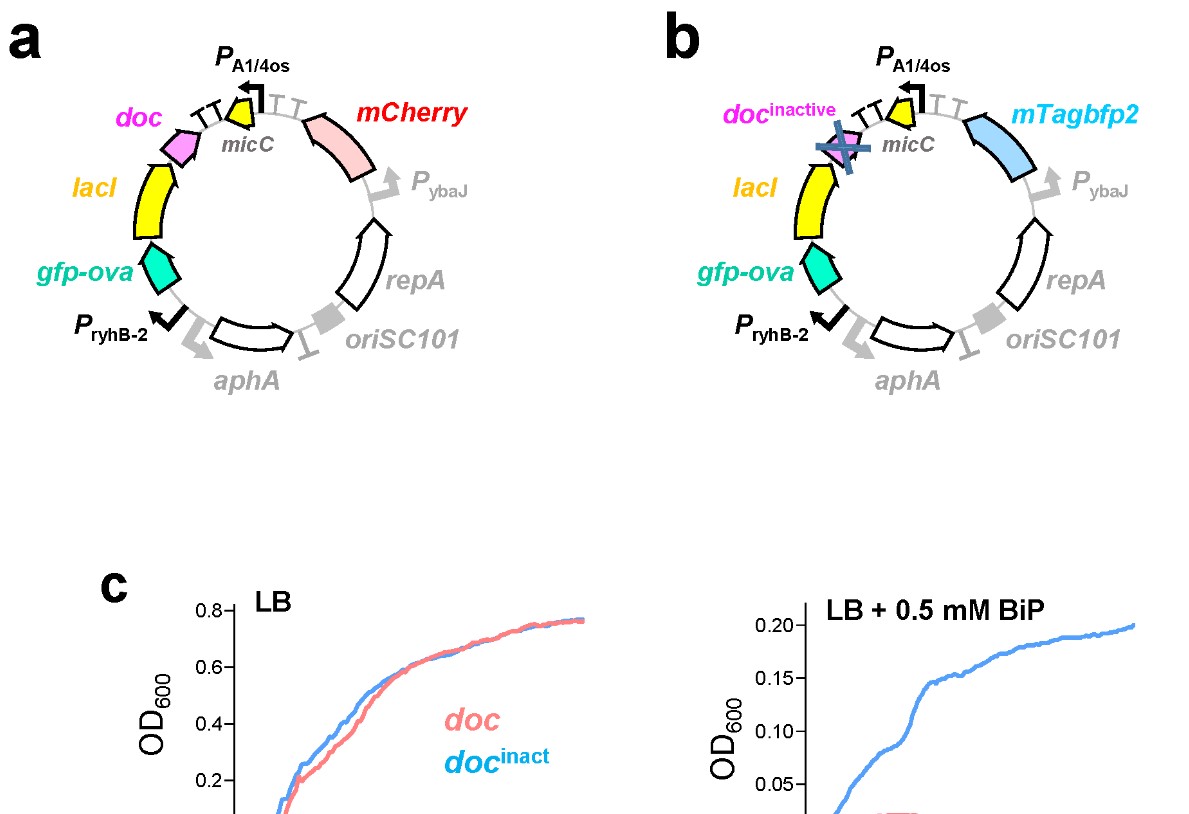

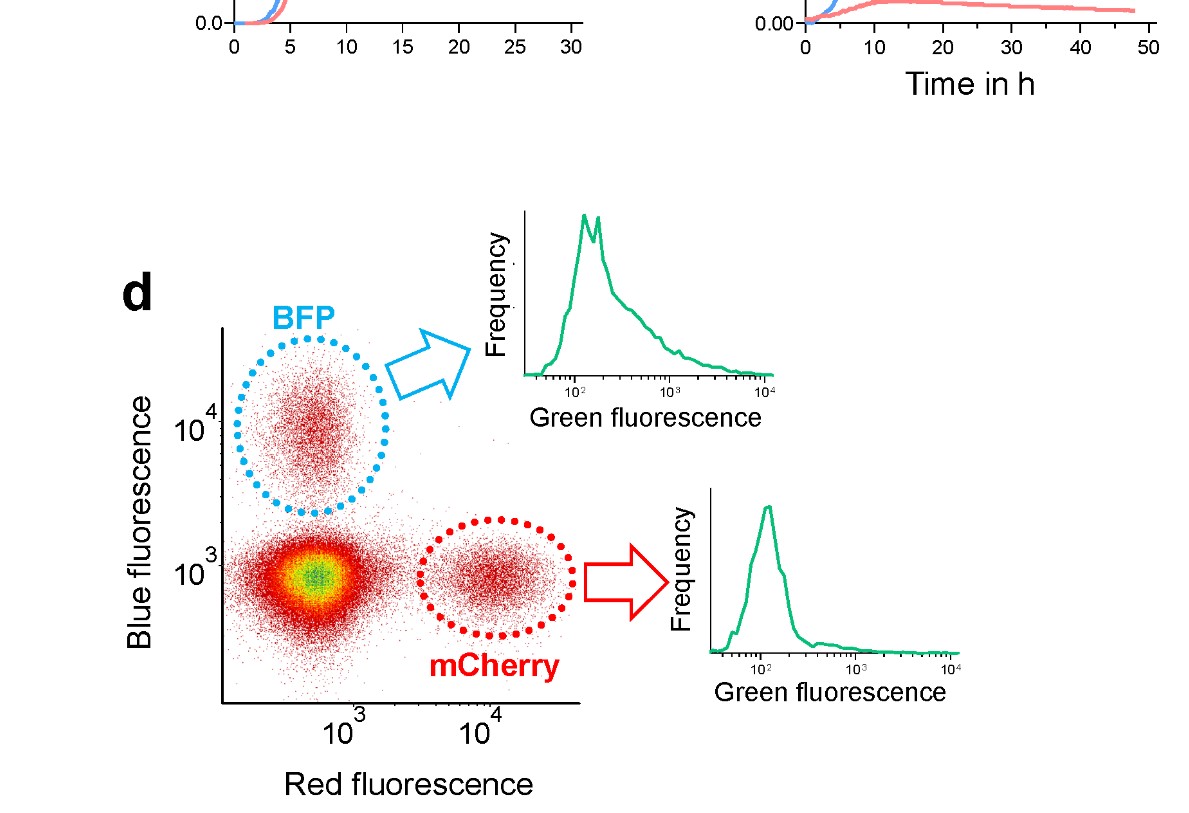

**fig. s7. Depletion of iron-starved *Salmonella*.**

(a) Detailed plasmid map for construct with conditionally expressed active toxin gene *doc*. For explanation of gene names see text and Fig. 3A; sfig. 3a.
(b) Detailed plasmid map for the control construct with inactive *doc*.
(c) Growth of *Salmonella* carrying the active doc construct or the inactive control construct in iron-replete (left) or iron-starved conditions. Similar results were obtained in two independent experiments.
(d) Gating strategy for detecting the test *Salmonella* strain with conditionally active doc and mCherry, and the control strain with inactive *doc* and BFP.

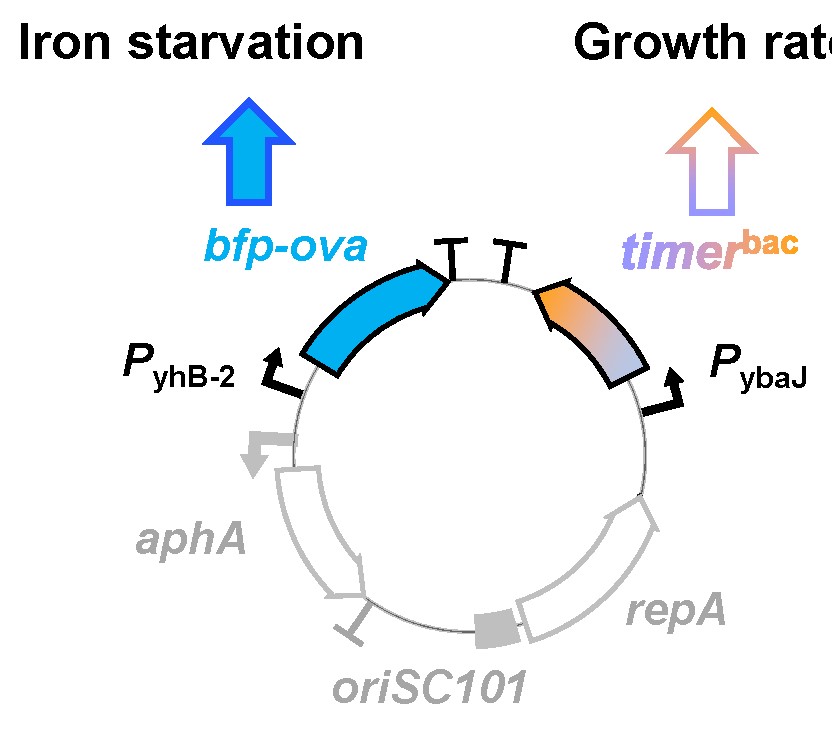

**fig. s8. Replication of *Salmonella* subsets with differential iron access.**

Plasmid map of a dual reporter construct for determining in individual *Salmonella* cells both both *P*_ryhB-2_*-bfp-ova* activity and division rate using *timer*^bac^ expressed from the constitutively active *P*_ybaJ_ promoter. For gene names see sfig. 3a.
